# Directed assembly of single-stranded DNA fragments for data storage via enzyme-free catalytic splint ligation

**DOI:** 10.1101/2024.10.09.617455

**Authors:** Gemma Mendonsa, Sriram Chari, Mengdi Bao, Brett Herdendorf, Anil Reddy

## Abstract

Oligonucleotides or gene fragments can be ligated in a specified order to create longer DNA assemblies. We present a method where DNA symbols, or oligos designed to encode information for data storage, are joined to linker sequences at either end. These linkers dictate the assembly order of the symbols; the order of the symbols can be changed by changing the sequences of the linkers attached to them. Utilizing a ligating DNAzyme as a catalytic splint, we achieve room-temperature, enzyme-free assembly, offering a cost-effective alternative to traditional enzyme-based ligation methods. We demonstrate this technique by assembling three different five-symbol constructs, with the order of the symbols determined by their linking ends. This linker directed assembly technique allows data-encoding symbols to be assembled in any desired order. Furthermore, the DNAzyme-based assembly method is versatile and can be applied to various DNA assembly applications, particularly where cost-effectiveness and efficient room-temperature ligation are required.

## INTRODUCTION

The growth of data generation, by both human and machine sources, is projected to reach 175 zettabytes (1ZB = 10^21^B) annually by 2025 (1). However, the fact that less than 0.5% of this data is archived on offline media like tape and optical disc underscores the challenges in data storage and preservation. Archiving digital data necessitates significant physical space and electrical power, depending on the storage technology employed. In addition, most storage systems, due to media durability or the obsolescence of writer/reader hardware, or of data formats or protocols, require periodic migration of archived data to fresh media (2). To be able to archive more of the newly generated data, paradigm-shifting new storage methods are a necessity.

DNA, nature’s medium for genomic data storage, offers an unprecedented level of volumetric storage density. A single gram of DNA can theoretically store up to 215 petabytes (1PB = 10^15^B) (3), which is several orders of magnitude greater than solid-state drives (SSD), hard disk drives (HDD), and tape media. (https://www.micron.com/about/blog/applications/data-center/dnas-awesome-potential-to-store-the-worlds-data). Furthermore, DNA has a half-life of approximately 500 years and can be replicated with relative ease, thereby reducing the need for new storage media solely for the purpose of refreshing previously archived data (2).

The transition of DNA data storage from a research concept to a product ecosystem under development, and subsequent broad usage, will require achieving several performance and cost measures. Write (molecular synthesis) rates and read (DNA sequencing) rates must be sufficiently great, and hardware and chemical reagent costs sufficiently low, that the overall cost of ownership to the customer over the life of the archive is competitive with incumbent archival technologies, such as tape.

DNA data storage typically relies on phosphoramidite chemistry, a well-established technology for synthesizing short DNA strands, one nucleotide at a time. The imperfect coupling between bases (coupling efficiency ≈ 0.99) limits the length of the synthesized oligos. For example, for a 200-mer oligo, the yield is only ≈13.4% (= 0.99^200^) (4). The coupling time for phosphoramidite chemistry is slow, around 4-10 min per base (5). Phosphoramidite-based methods also require organic solvents, resulting in hazardous waste (6). The conservative estimate for the amount of data expected to be stored by 2040 is ≈ 10^24^ bits, with certain estimates being as high as ≈ 10^24^ bits (4)(7). Based on these projections, Baek et al. (4) estimated the total cost to write this data in DNA using phosphoramidite chemistry to be ≈ $10^23^ - $10^25^. Even if 1% of this data is archived, the cost can be as high as ≈ $10^21^ - $10^23^, which makes it a prohibitively expensive proposition. Consequently, this synthesis process has become a significant bottleneck in the development of DNA storage technologies. Enzymatic DNA synthesis can address some of the limitations of phosphoramidite chemistry by using more environmentally friendly aqueous solvents to synthesize longer DNA strands with potentially higher coupling rates (8)(4). Different enzymatic approaches for synthesizing and assembling oligonucleotides for DNA-based data storage have been reviewed in literature (9)(4). Most of the popular enzymatic synthesis technologies involve template independent enzymatic oligonucleotide synthesis (TIEOS), which uses the enzyme terminal deoxynucleotidyl transferase (TdT) for synthesis (10). However, these approaches have their unique set of limitations. For example, in TIEOS, TdT tends to favor attachment of certain nucleotides over others, which may lead to sequence-specific errors (11)(9). The main challenge for enzymatic synthesis technologies remains throughput. With phosphoramidite chemistry, research groups and companies have achieved high throughput by using a microarray-based platform for synthesizing the storage oligo. Progress has been made in the area of parallel enzymatic DNA synthesis in recent years; however, a commercially viable and practical technology is still not mature enough for practical DNA data storage (12).

To achieve practical write speeds, three key elements are needed: a chemical protocol that maximizes the data written per operation through parallelization and a method which is cost-effective and environmentally friendly. To resolve the speed of synthesis problem, oligo assembly-based approaches have been proposed in literature. Yan et al. (13) introduced an enzymatic ligation technique for synthesizing storage genes based on composite motifs as building blocks. A dual-library system of “symbols” and “linkers” has also been proposed (14)(15). The “symbols” carry the data payload and the “linkers” are short oligos that help concatenate these symbols in the correct order. This approach alleviates the need to build storage oligos sequentially, one nucleotide at a time. These libraries work together to create long “genes” with symbols in the correct order, representing the data. By using linkers, a ‘one-pot’ reaction to build the desired storage oligo can be realized, enabling massive parallelization. One aim of this work is to demonstrate this dual-library assembly approach by assembling symbols in specified orders to create several different data storage genes.

A major drawback of any enzymatic process, whether it be oligo synthesis or assembly, is the cost of the enzymes. While the costs of protein-based enzymes are manageable at benchtop-scale experiments, when the reagent volumes are scaled up to data-storage level volumes the cost becomes exorbitant problematic. Synthesizing oligos in bulk is a much cheaper process than synthesizing each individual data storage gene from scratch (13), but the high cost of the ligase used to ligate the oligos still results in inordinately expensive DNA assembly. For example using the current pricing of DNA assembly reagents from NEB, it is estimated that the total cost of the reagents needed for ligating oligos together using standard T4 ligase is $1.36 for a 20µL reaction (https://www.neb.com/en-us/products/m0202-t4-dna-ligase), while the cost of the ligase is $1.35 (https://www.neb.com/en-us/products/b0202-t4-dna-ligase-reaction-buffer), making up the majority of the cost. While the oligo assembly-based approaches described above can make DNA synthesis faster, the cost of these methods remains prohibitively high for DNA data storage use cases

The second aim of this work is to introduce a novel cost-effective DNA assembly method that requires no protein-based enzymes. Our approach involves the use of DNAzymes for concatenating short DNA segments through chemical ligation to build the desired storage gene (16). The DNAzymes act as catalytic molecular splints, bringing together the ends of DNA strands and catalyzing the formation of phosphodiester bonds. This process eliminates the need for ligase or any other protein-based enzymes during the ligation step (17). DNA ligation using DNAzymes uses aqueous solvents, which lowers its environmental footprint (4). We describe here a synthesis strategy and ligation chemistry that has the potential to offer the DNA data storage community a means to achieve geometric DNA strand growth, rather than the nucleotide-by-nucleotide synthesis strategy employed by many researchers in this field, in a cost-effective and environmentally friendly manner.

## MATERIALS AND METHODS

### Materials

The oligos used in the Figure 2 experiment were synthesized using an H-8 oligo synthesizer by K&A Laboratories according to the manufacturer’s instructions. Size markers used in the gel electrophoresis experiments in Figure 4b and Figure 5b were synthesized by GENEWIZ from Azenta Life Sciences. All other oligos and ssDNA ladders were purchased from Integrated DNA Technologies. Phusion PCR kit, dNTPs, Zeba spin columns, TBE buffer, HPLC-grade acetonitrile, PureLink Gel Extraction kit, and 100bp ladder were purchased from Thermo Scientific. Certified Low-range agarose was purchased from BioRad. DeepVent and DeepVent (exo-) polymerases and PCR buffers were purchased from NEB. SD Polymerase was purchased from Bioron. NAP-5 desalting columns were purchased from Cytiva. Sanger Sequencing reagents were purchased from Promega. Imidazole, EDC, ZnCl, HEPES, NaCl, PAGE gel electrophoresis reagents, Triethylammonium acetate, and SYBR Green I/II gel stains were purchased from Sigma Aldrich. DNA sequences are available in the Supplementary Information (Supplemental Table 1).

### 2-piece DNAzyme Activation and Ligation

#### S1 Subunit Activation

Fresh 1M EDC solution was prepared. A solution of 80µM S1 subunit, 100mM, 20mM, or 5mM imidazole (pH 6.0), and 100mM EDC was mixed, vortexed, and incubated on the benchtop at room temperature for 60min. If a purification step was being performed, the activation solution was desalted with a Zeba 7K MWCO spin column according to manufacturer’s instructions. The DNA amount remaining in the desalted solution was quantified by a NanoDrop 1C. Activated S1 was immediately used in a ligation reaction.

#### 2-piece ligation

A solution of 2µM S2 subunit, 3µM catalytic strand, 4µM activated S1 subunit, and 4mM ZnCl was prepared in a buffer containing 30mM HEPES and 300mM NaCl at pH 7.0. Negative controls had no ZnCl added. The ligation solutions were incubated at room temperature on the benchtop for 60min. After the ligation period a 1/10 volume of 100mM EDTA (pH 8.0) was added to the ligation solutions.

### 3-piece DNAzyme Activation and Ligation

#### S1 Subunit Activation

Fresh 1M EDC solution was prepared. A solution of 80µM S1 subunit, 20mM imidazole (pH 6.0), and 100mM EDC was mixed, vortexed, and incubated on the benchtop at room temperature for 60min. The activation solution was desalted with a Zeba 7K MWCO spin column according to manufacturer’s instructions. The DNA amount remaining in the desalted solution was quantified by a NanoDrop 1C. Desalted activated S1 was immediately used in a ligation reaction.

#### 3-piece Ligation

A solution of 2µM S2 subunit, 4µM catalytic strand A, 4µM catalytic strand B, 4µM activated S1 subunit A, 4µM activated S1 subunit B, and 4mM ZnCl was prepared in a buffer containing 30mM HEPES and 300mM NaCl at pH 7.0. Negative controls had no ZnCl added. The ligation solutions were incubated at room temperature on the benchtop for 120min. After the ligation period a 1/10 volume of 100mM EDTA (pH 8.0) was added to the ligation solutions. The full-length ligation products were purified by HPLC and desalted with NAP-5 columns according to the manufacturer’s instructions. The desalted ligation product was concentrated in a 60°C vacuum centrifuge for 1.5hrs.

### 5-piece DNAzyme Activation and Ligation

#### S1 Subunit Activation

Fresh 1M EDC solution was prepared. A solution of ≈ 1µM of each of four linker-symbol assemblies, 20mM imidazole (pH 6.0), and 100mM EDC was mixed, vortexed, and incubated on the benchtop at room temperature for 60min. The activation solution was not desalted, but immediately used in a ligation reaction.

#### 5-piece Ligation

A solution of 0.1µM S2 subunit, 0.1µM each of 4 catalytic strands, and ≈0.4µM activated S1 subunit mix, and 4mM ZnCl was prepared in a buffer containing 30mM HEPES and 300mM NaCl at pH 7.0. Negative controls had no ZnCl added. The ligation solutions were incubated at room temperature on the benchtop for 3hrs. After the ligation period a 1/10 volume of 100mM EDTA (pH 8.0) was added to the ligation solutions. The full-length ligation products were analysed and purified by gel electrophoresis.

### High Performance Liquid Chromatography (HPLC) Analysis and Purification

HPLC was performed on an Agilent 1100 series HPLC system with DAD detector and fraction collector. The Waters XBridge Oligonucleotide C18 BEH column was heated to 80°C. Mobile Phase A consisted of 100mM Triethylammonium Acetate (TEAA) buffer (pH7.0). Mobile Phase B consisted of 80mM TEAA and 20% v/v HPLC-grade acetonitrile. A gradient of 60%A / 40%B was adjusted to 40.5%A / 59.5% B over the course of 7.5min at 0.6ml/min. If purification was being performed, ligation products were collected by a fraction collector and concentrated in a 60°C vacuum centrifuge for 30min. Standards of the known ligation product were run, and the resulting peak areas plotted vs concentration with JMP. Standard curves, equations, and fits were generated using JMP (Supplementary Figure S2A-2C).

### Gel Electrophoresis (PAGE)

A freshly prepared 1.6% m/v solution of ammonium persulfate (APS) was prepared in ultrapure water. A gel solution of 12% acrylamide and 8M urea was prepared in 1X TBE. 2.2mL of the APS solution was added for every 50mL of gel. 33.3µL of TEMED was added for every 50mL of gel, and the gel was immediately cast in 20cm *×* 20cm plates with separated by 1.5mm spacers. The gel polymerized for 90min, then was loaded into a BioRad Protean Xi electrophoresis core. Chilled water was run through the core while the gel pre-ran at 300V for 30min. Samples were mixed with formamide in a 1:1 volume ratio, heated to 80°C for 5min, then plunged into an ice bath for 5min before loading. The gel was run at 450V for 4hr with chilled water running through the core. The gel was removed from the plates and stained in 1X SYBR Green II for 20min. The gel was rinsed 3*×* with ultrapure water and imaged in a GelDoc Go imager by BioRad Laboratories. Desired bands were excised, crushed with a plastic dowel, and soaked in molecular grade water on a rotator for 24hr. The gel extract was filtered with a 0.22µm syringe filter, concentrated in a 60°C vacuum centrifuge for 30min, and desalted with NAP-5 columns according to the manufacturer’s instructions.

### Polymerase Chain Reaction (PCR)

DNA was amplified using 1X reaction buffer (from various kits, the reaction buffer always was paired with the polymerase used), 200µM dNTPs, 300nM each forward and reverse primers, and 0.02 – 0.04U/µL polymerase. If ET SSB was used, it was added to a concentration of 4ng/µL. For amplifications with Phusion, DeepVent, and DeepVent (exo-), the T100 thermocycler from BioRad was set to perform an initial denaturation step at 95°C for 2min, followed by 30 cycles of 95°C for 30s, 58°C for 30s, and 72°C for 15s. A final extension at 72°C was performed for 5min followed by a 4°C hold. For the amplifications with SD Polymerase, 3mM MgCl2 was added and the thermocycler program set to perform an initial denaturation step at 92°C for 2min, followed by 30 cycles of 92°C for 30s, 58°C for 30s, and 68°C for 15s. A final extension at 68°C was performed for 5min followed by a 4°C hold.

### Agarose Gel Electrophoresis

PCR products were run on a 2% agarose gel in 1X TBE. 100bp ladder was loaded into the first well. The gel was run at 95°C for 45min. Any desired bands were excised and purified with the PureLink Quick Gel Extraction Kit from Invitrogen.

### Sanger Sequencing

#### Dideoxynucleotide PCR

PCR products were sequenced using the ProDye sequencing system according to the manufacturer’s instructions. Approximately 50ng PCR product was mixed with 1X ProDye Master Mix and 5pmol of either the forward or reverse primer. The thermocycler program was set to perform an initial denaturation at 96°C for 1min, followed by 25 cycles of 96°C for 10s, 50°C for 5s, and 60°C for 4min.

#### Ethanol Precipitation

Dideoxynucleotide PCR products were cleaned up by ethanol+EDTA precipitation. 5µL of 125mM EDTA (pH8.0) and 60uL 100% ethanol were added to each 20µL sequencing reaction. Samples precipitated at room temperature for 15min, then centrifuged at 14000RCF in a chilled centrifuge for 15min. The supernatant was pipetted off and the pellet washed with 70% ethanol at 14000RCF for 5min. Supernatant was removed and pellets dried in a vacuum centrifuge for 5min. The pellets were resuspended in 10µL HiDye formamide.

#### Sequencing

Sequencing was performed on the Spectrum Compact by Promega using Polymer 7. Data analysis. Sequencing data was analysed using SnapGene.

#### Data Analysis

Sequencing data was analyzed using SnapGene.

## RESULTS

### DNAzyme characterization

The E47 DNAzyme (18) was selected as the best candidate for ligating two DNA strands end-to-end after searching the DNAzyme database DNAmoreDB (19). Several ligating DNAzymes exist, but the majority of the ones in the database form a branched DNA product (19)(20)(21). In other cases, the ligating DNAzymes are long, such that synthesis becomes more arduous and expensive (18) or are highly sequence dependent (22). The catalytic region of the E47 DNAzyme is only 24nt long, with 10nt and 13nt long binding arms for hybridizing the two subunit DNA strands together (Figure 1a). A 5’ OH group is required on the S2 subunit, and a 3’ phosphate group is required on the S1 subunit. The 3’ phosphate is activated with imidazole and EDC to generate a reactive phosphorimidazolide. Then the activated S1 unit is purified and mixed with the S2 subunit, the E47 catalytic strand, and Zn2+ as a cofactor. Only four nucleotides of the subunits to be ligated interact with the catalytic region of the E47 DNAzyme; it seemed reasonable to explore whether the binding arm sequences of the E47 DNAzyme could be changed as long as the four nucleotides in the catalytic region remained.

**Figure 1.**
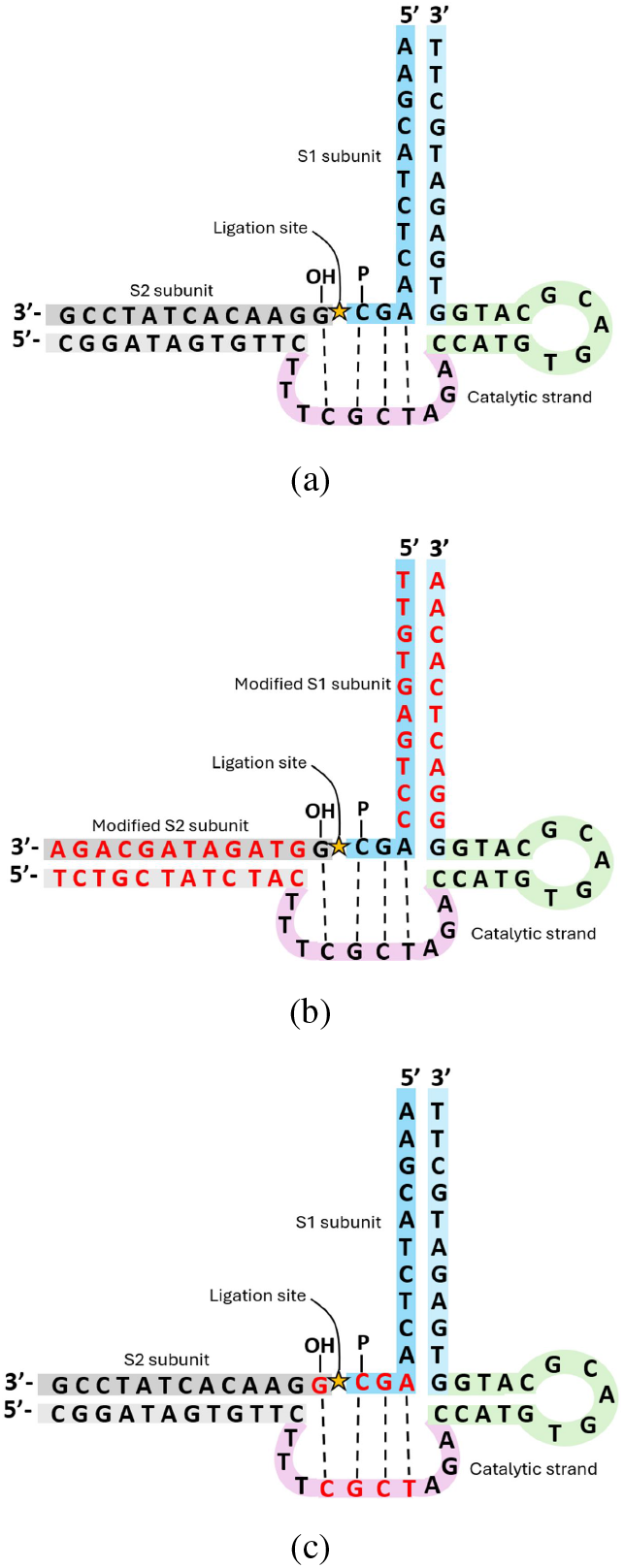
(a)The original E47 DNAzyme and subunits as reported by Cuenoud and Szostak (18). **(b)** E47 with modified substrates and substrate binding arms. **(c)** E47 4-base catalytic motif. The bases in this region are likely directly involved in catalysis. The highlighted bases (red text) were mutated one by one to explore the sequence dependence of this region.

Sequence specificity of the E47 DNAzyme. The 5’ G on the S2 strand and the last three nucleotides on the 3’ end of the S1 strand seem to interact with the catalytic region of the DNAzyme. However, the other nucleotides in the substrate strands seem to be mostly involved in hybridization to the DNAzyme, and may not play a role in the catalysis. We explored the sequence specificity of the E47 DNAzyme by changing the binding arm sequence of the substrates and DNAzyme to random sequences (Figure 1b). The resulting ligation products were analyzed with HPLC. The ligation product peaks were strong in both samples, indicating that the binding arm regions of the E47 DNAzyme can tolerate sequence modifications (Figure 2a - Figure 2b).

**Figure 2.**
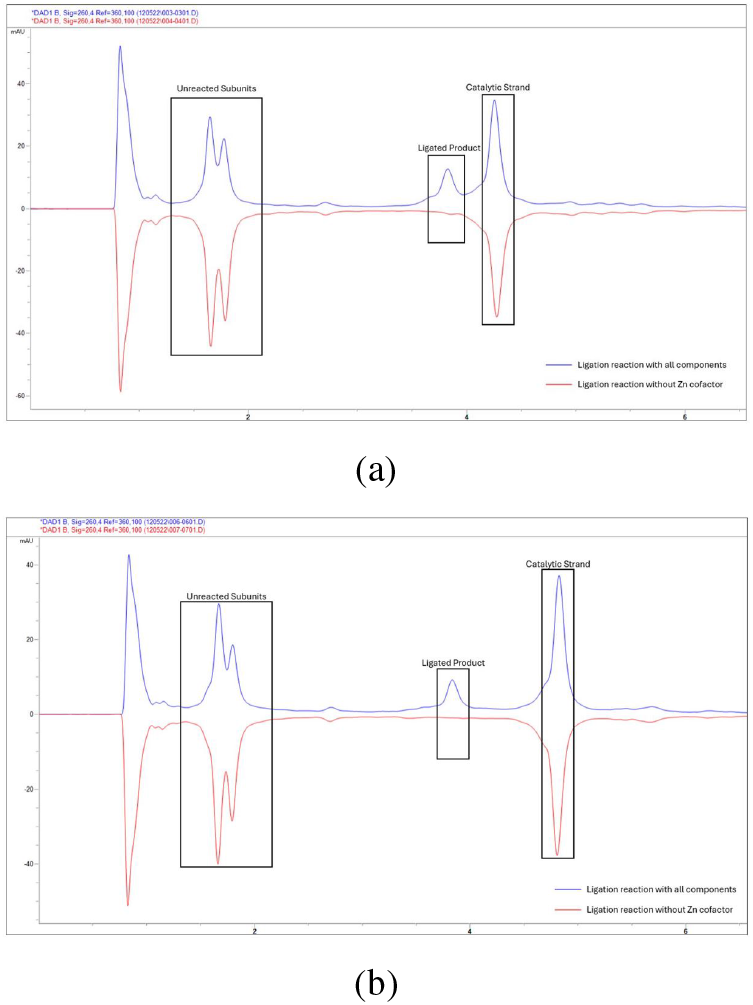
(a)HPLC chromatograms of ligation products with the original reported sequence. Top (blue) line: Ligation reaction with all components. Bottom (red) line: Ligation reaction without Zn cofactor. **(b)** HPLC chromatograms of ligation products with modified subunit sequences. Top (blue) line: Ligation reaction with all components. Bottom (red) line: Ligation reaction without Zn cofactor.

The sequence specificity of the catalytic 4nt motif was also probed (Figure 1c). Each nucleotide was either changed to its complement, maintaining the base pair identity in that position ( i.e. G → C or T → A), or switched to the other nucleotide that maintained the pyrimidine/purine identity in that position (i.e. C → T or G → A). Any of these modifications to the 4nt catalytic motif resulted in a complete loss of ligation activity as measured with HPLC, with the exception of the 3’ T in the catalytic strand. Modifications of the 3’ T resulted in reduced catalytic activity and unexpected ligation products (Table 1, Supplementary information S1 A-C). While the E47 DNAzyme can ligate a variety of sequences, the S1 strand must have the motif AGC at its 3’ end, while the S2 strand must have a G in the 5’ end position.

**Table 1.**
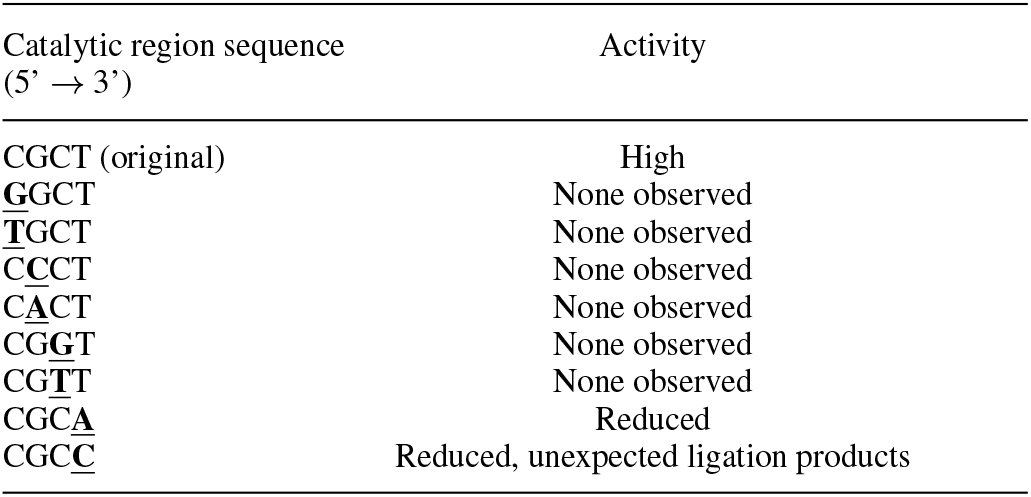
Ligation activity of catalytic 4-base motif mutations.

#### Necessity of purification step between activation and ligation

There are two steps in performing ligation with this DNAzyme: activation of the 3’ phosphate group with imidazole and EDC, and ligation. Historically, a purification step was included between the activation and ligation steps to remove unreacted EDC and imidazole. The purification step is time consuming, sometimes leading to inactivation of the unstable phosphorimidazolide before the ligation could take place. The purification step also adds cost of purification materials such as columns, and results in the loss of some subunit due to column retention. We explored removing this step as well as the possibility of combining the activation and purification steps in a one-step reaction.

Purification between the activation and ligation steps is desired if one or both of the activation reactants (EDC or Imidazole) inhibits the subsequent ligation reaction. To probe the inhibitory nature of EDC and imidazole, ligation reactions were spiked with either EDC, imidazole, or EDC + imidazole. Results show that while spiking the ligation reaction with EDC did not have a notable effect on the ligation reaction, imidazole showed a strong inhibitory effect (Figure 3a). Next, ligation reactions were attempted without the use of a purification step between the activation and ligation steps, with varying amounts of imidazole in the activation step. Reducing the imidazole concentration by from 100mM to 20mM in the activation reaction still showed good activation of the substrate, and removal of the purification step only minimally reduced the ligation activity (Figure 3b). Combining the activation and ligation steps resulted in a marked reduction in ligation activity (Figure 3b and Figure 3c), even with reduced imidazole concentrations. The activation and ligation reactions could therefore not practically be combined in a one-pot reaction, but if desired, the purification step between the activation and ligation steps can be removed.

**Figure 3.**
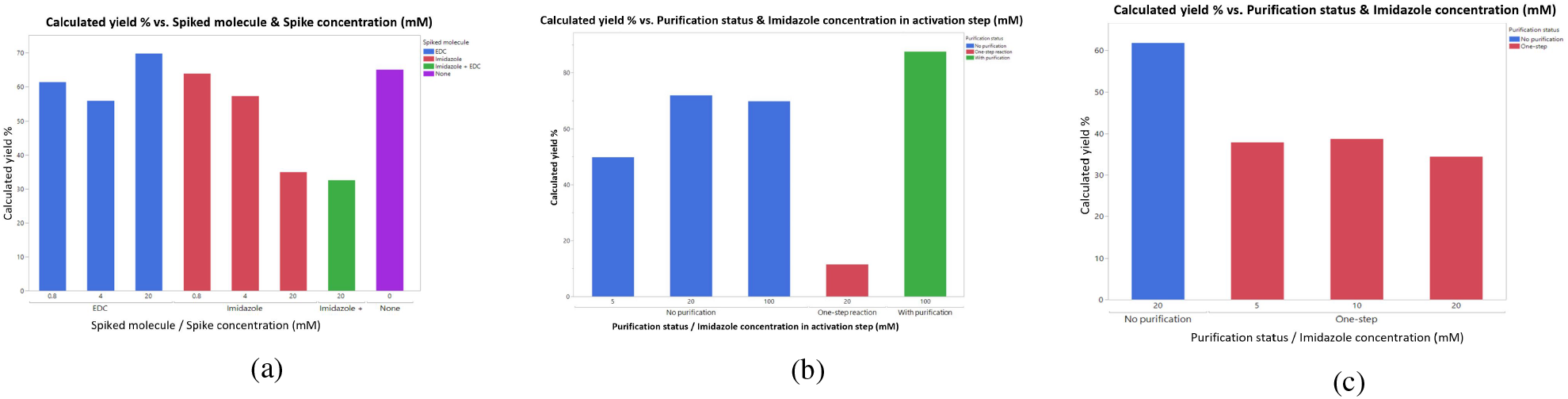
(a)Varying amounts of EDC, imidazole, or both EDC and imidazole were spiked into the ligation reaction. The final concentrations of EDC and imidazole after spiking the ligation solutions were plotted vs. calculated yield as quantified by HPLC. **(b)** The purification step between activation and ligation steps was removed, and/or the activation and ligation steps were combined in a one-step reaction. Varying levels of imidazole were used to activate the S1 substrate. **(c)** The ligation and reaction steps were combined in a one-step reaction and the amount imidazole reduced.

### First-tier ligation: Attaching linkers to the symbols

The first step in assembling the DNA data storage gene in our schema is to attach unique linkers to each symbol. Each symbol was one of three sequences, designated as Symbols A, B, or C. The linkers attached to each symbol were selected based on the desired order of the symbols in the final assembly. For example, in order to have the symbols assembled in the order A → B → C → B → A, the symbols would have to be matched to the linkers that would attach them in that order (Table 2). Each linker possesses two regions: one region that connects to the symbol, and one that connects to another linker (Figure 4a). The symbol-connecting region is the same for each linker, allowing any linker to be attached to any symbol. Conversely, the linker-connecting region is different for each, allowing only two specific linkers to be joined to each other. Sequences of each strand can be found in the Supplementary information. The linker-symbol assembly requires a three-piece ligation reaction involving three substrate strands and two unique DNAzymes (Figure 4a). Analysis of the products with gel electrophoresis showed that the three-piece ligation was successful (Figure 4b), and the full-length product was purified from the reactants using either HPLC or gel electrophoresis. Five different linker-symbol complexes were prepared in this manner. At this stage PCR was not performed, because for the next ligation reaction the pieces need to have a 3’ phosphate. PCR amplification only results in 3’ OH groups. In addition, PCR results in double-stranded DNA, whereas for the next ligation reaction, single-stranded DNA is required.

**Figure 4.**
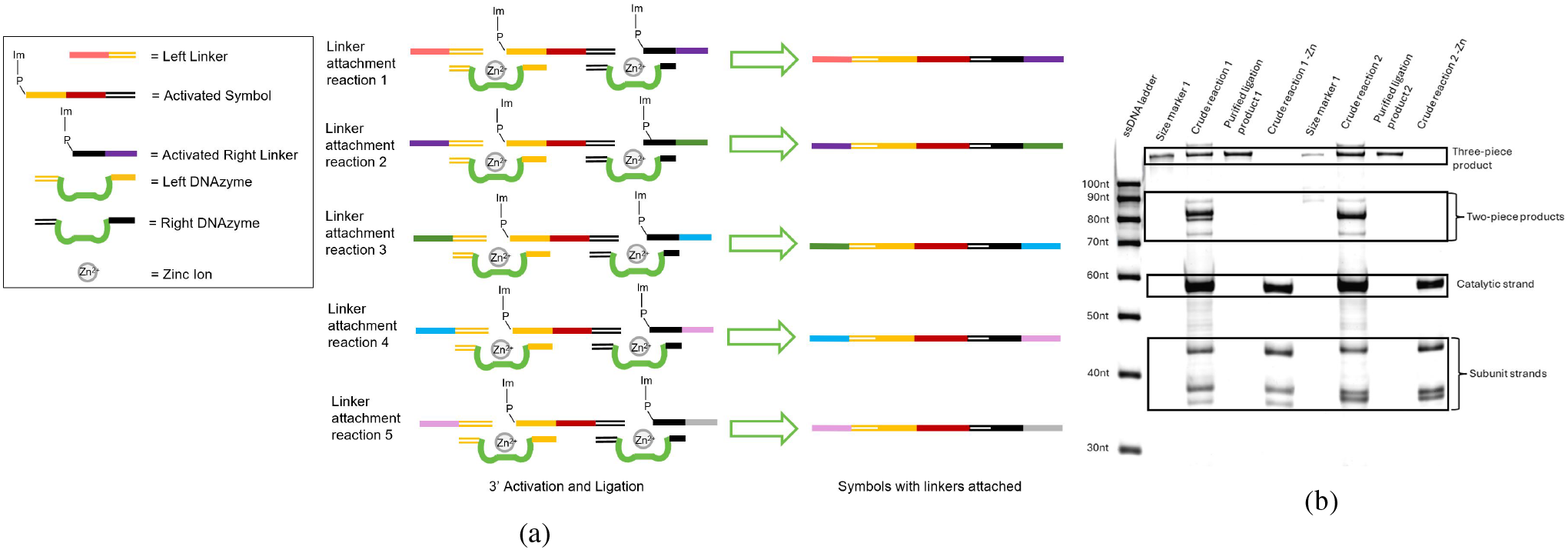
(a)Schematic of three-piece ligation and gel image of three-piece assembly. **(b)** Gel image of two different three-piece assemblies, including size markers, crude reaction, purified ligation product, and crude reaction with no Zn cofactor.

**Table 2.** The order of the data-encoding symbols was directed via the linker sequences. The order was changed by either changing the symbol-linker pairings in the first assembly step as for Storage Gene 2, or by changing the catalytic strand sequences in the second assembly step as for Storage Gene 3.

| Symbol order in the final data storage gene 5' → 3' | Linkers added to each symbol in assembly step 1 | Catalytic strands used to join the linkers in assembly step 2 |
| --- | --- | --- |
| Data Gene 1:<br>A, B, C, B, A | Symbol A, Left Linker 1, Right Linker 1<br>Symbol B, Left Linker 2, Right Linker 2<br>Symbol C, Left Linker 3, Right Linker 3<br>Symbol B, Left Linker 4, Right Linker 4<br>Symbol A, Left Linker 5, Right Linker 5 | Joins Right Linker 1 to Left Linker 2<br>Joins Right Linker 2 to Left Linker 3<br>Joins Right Linker 3 to Left Linker 4<br>Joins Right Linker 4 to Left Linker 5<br>N/A |
| Data Gene 2:<br>B, C, A, C, B | Symbol B, Left Linker 1, Right Linker 1<br>Symbol C, Left Linker 2, Right Linker 2<br>Symbol A, Left Linker 3, Right Linker 3<br>Symbol C, Left Linker 4, Right Linker 4<br>Symbol B, Left Linker 5, Right Linker 5 | Joins Right Linker 1 to Left Linker 2<br>Joins Right Linker 2 to Left Linker 3<br>Joins Right Linker 3 to Left Linker 4<br>Joins Right Linker 4 to Left Linker 5<br>N/A |
| Data Gene 3:<br>C, B, C, A, B | Symbol B, Left Linker 1, Right Linker 1<br>Symbol C, Left Linker 2, Right Linker 2<br>Symbol A, Left Linker 3, Right Linker 3<br>Symbol C, Left Linker 4, Right Linker 4<br>Symbol B, Left Linker 5, Right Linker 5 | Joins Right Linker 2 to Left Linker 1<br>Joins Right Linker 1 to Left Linker 4<br>Joins Right Linker 4 to Left Linker 3<br>Joins Right Linker 3 to Left Linker 5<br>N/A |

### Second-tier ligation: Assembling the data symbols

#### Five-piece DNA assembly

The five symbol-linker complexes were assembled in a second ligation reaction, with four DNAzymes designed to hybridize to the linker ends and join them in a specified order (Figure 5a), resulting in a five-symbol data storage gene. The results were verified by gel electrophoresis and the full-length product purified from the gel (Figure 5b). The five-piece product was amplified by PCR and the product purified via an agarose gel.

**Figure 5.**
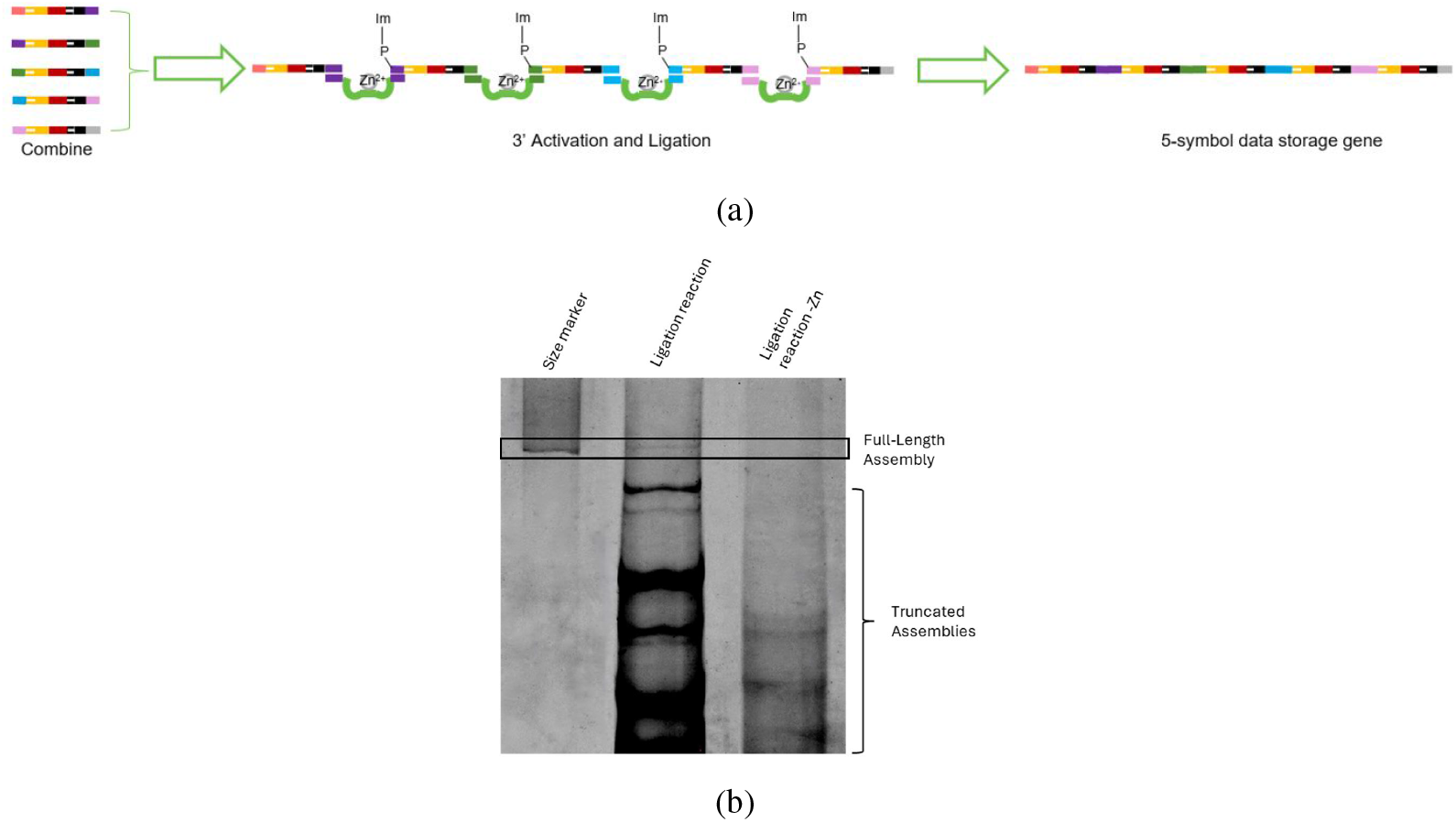
(a)5 symbols with linkers attached were assembled in a final ligation reaction to create a 5-symbol data storage gene. **(b)** Final assembly of five sub-assemblies. Lane 1: Size marker. Lane 2: Ligation reaction. Lane 3: Ligation reaction with no Zn cofactor.

#### PCR optimization of the assembled DNA data storage gene

PCR of the five-piece assembly with Phusion polymerase results in a ladder banding pattern due to the repeats of the linker attachment sequence in the template (Figure 6a) (23). Changing to a different DNA polymerase with strand displacement activity may result in a reduction of the ladder banding (23). Several different DNA polymerases were probed, in addition to a single-stranded DNA binding protein (ET SSB) that is known to reduce secondary structure formation in difficult PCR templates (23). ET SSB addition had positive impacts on amplification with Phusion and SD HotStart, but a negative impact on amplification with DeepVent (exo-). Phusion polymerase and SD Polymerase showed strong amplification but lots of laddering (Figure 6A). DeepVent polymerase didn’t show any amplification, but a DeepVent polymerase modified to remove the 3’→ 5’ exonuclease showed good amplification with marked reduction in laddering (Figure 6a - Figure 6b). Three unique assemblies were prepared and amplified with DeepVent exo-polymerase, resulting in strong amplification of the full-length assembly and minimal laddering (Figure 6b).

**Figure 6.**
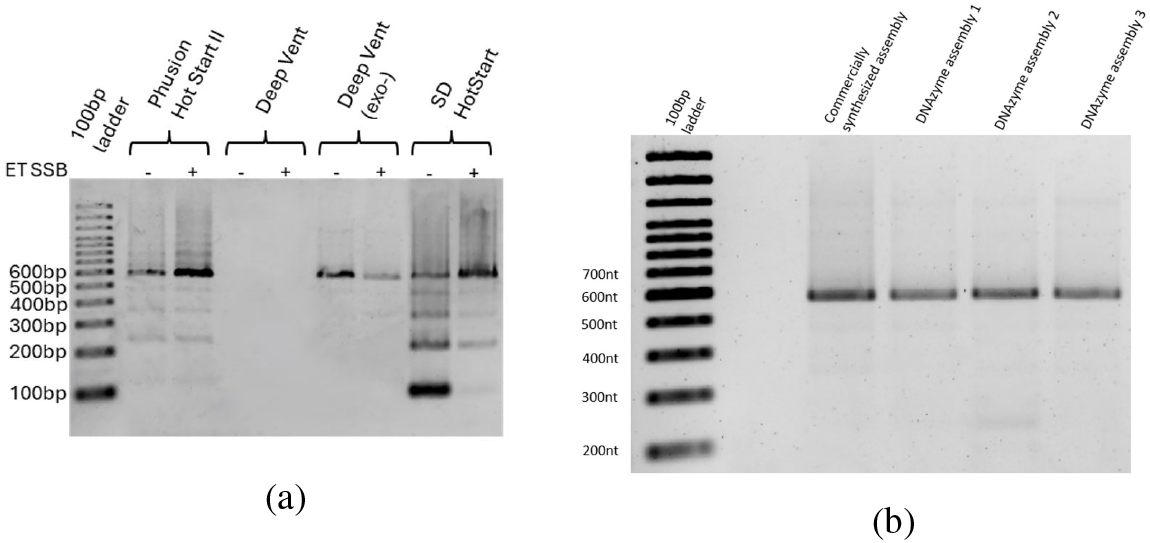
(a)PCR of a five-symbol data storage gene assembly with different polymerases with and without ET SSB. Expected gene length is 583bp. **(b)** PCR amplification of three data storage genes assembled with DNAzymes with DeepVent (exo-) polymerase.

#### Order of the data storage symbols directed via the linker ends

The order of the data-encoding DNA symbols can be directed by changing the linker ends attached to each symbol. Two data storage genes were prepared, and the order of the symbols was directed by switching the symbol-linker pairing in the first-tier assembly (Table 2). The symbol-linker complexes were assembled in a second-tier assembly reaction, and the resulting full-length product was purified and amplified as before. The two five-symbol data storage genes were sequenced and the correct symbol order verified (Figure 7a - Figure 7b).

**Figure 7.**
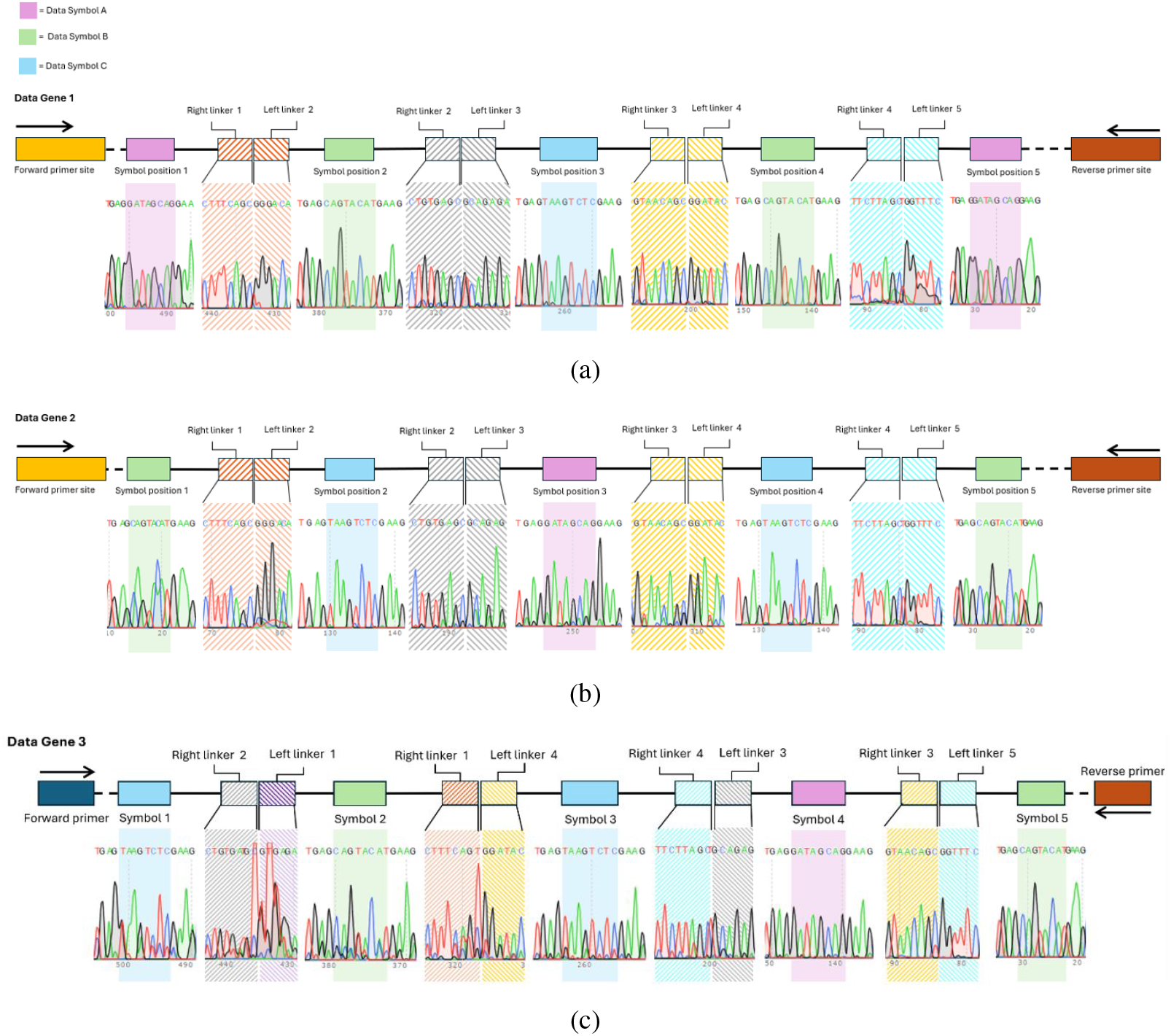
(a)Sequencing data with symbol and transition areas highlighted. A: Data Gene 1. **(b)** Data Gene 2 with symbols arranged in a different order via the Linker sequences. **(c)** Data Gene 3 with symbol+linker pieces arranged in a different order via the DNAzyme sequences.

A third five-symbol data storage gene was assembled, but this time the order of the symbols was shuffled by changing the sequences of the DNAzymes used in the second assembly step. The symbol-linker complexes used for this gene were the same as for the second gene, but the DNAzymes were designed to link them in a different order (Table 2). The sequencing data in Figure 7c shows that the third data gene was successfully assembled by changing the sequences of the DNAzyme catalytic strands in the second assembly step.

## DISCUSSION

We have demonstrated a novel DNA assembly method that can be leveraged for DNA data storage. The protein-free catalytic splint assembly approach reduces the cost of the assembly reagents, and the symbol-linker approach enables the use of bulk DNA oligos, which reduces the cost associated with synthesizing data storage oligos on demand (13). This chemical ligation method has other unique advantages; for example unlike other chemical ligation methods such as click chemistry, which introduce unnatural linkages in the DNA backbone, DNAzyme ligation preserves the natural phosphodiester backbone of DNA (24). The stability of the triazole linkages formed during click-ligation during long-term storage has not yet been explored, nor has their ability to be acted on by restriction endonucleases or the CRISPR/Cas editing system, which may be used in compute or editing operations. In addition, the catalytic splint ligation method only requires one oligo to have a modification (3’ phosphate), while click ligation requires modifications on both oligos (azide on one oligo, alkyne on the other) (25). Catalytic splint ligation could potentially be used for gene assembly for genomic applications; however, the necessity of the 4-base motif that interacts with the catalytic region of the splint does limit its versatility. Future work could explore novel ligating DNAzymes that do not possess such stringent sequence dependence in the catalytic region. In addition, exploration of novel ligating DNAzymes could also result in new DNAzymes that are not inhibited by imidazole, or that have a faster activity or higher yield than E47. Catalytic splint ligation is a promising avenue for DNA assembly, and particularly enables lower cost DNA data storage. Utilizing this chemistry paired with the joining of prefabricated data encoding symbols via linker ends makes fast, lower cost DNA data writing more feasible.

## DATA AVAILABILITY

The data underlying this article are available in the article and in its online supplementary material.

## SUPPLEMENTARY DATA

Supplementary Data are available at NAR online.

