## Supplementary Information for "Directed assembly of single-stranded DNA fragments for data storage via enzyme-free catalytic splint ligation"

### S1A.

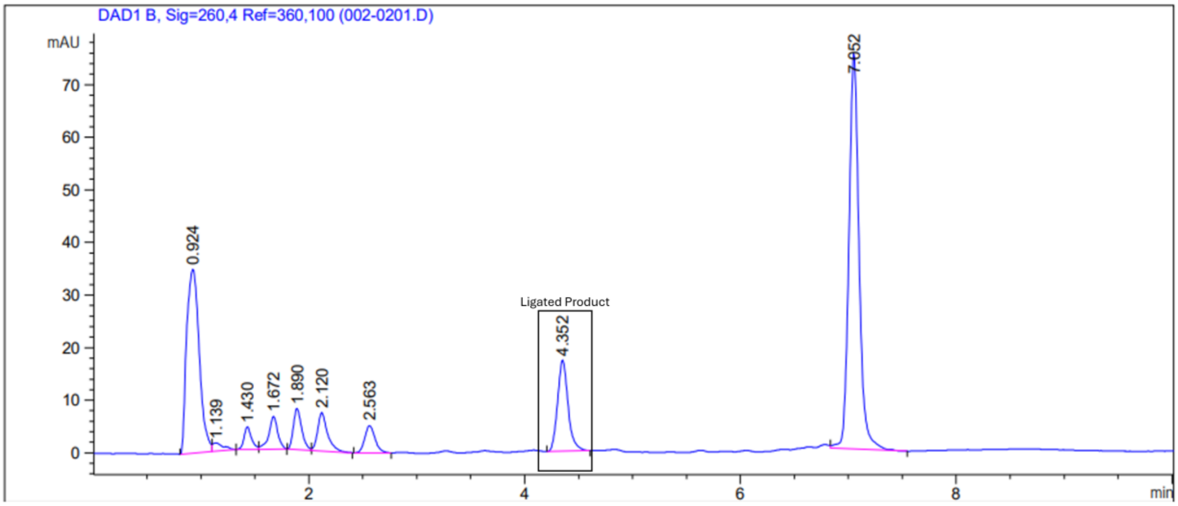


### S1B.

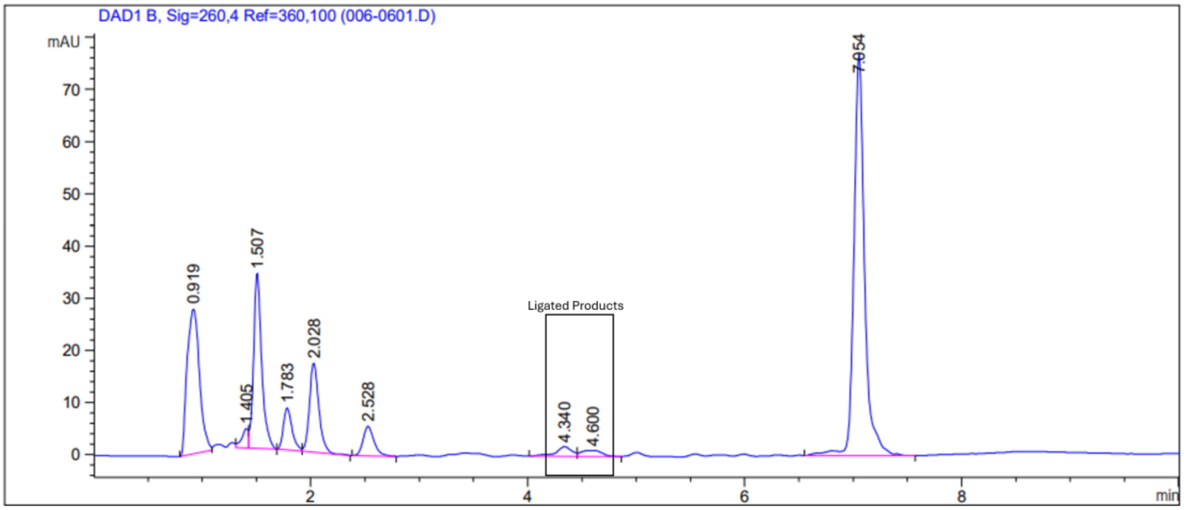

**S1C.**

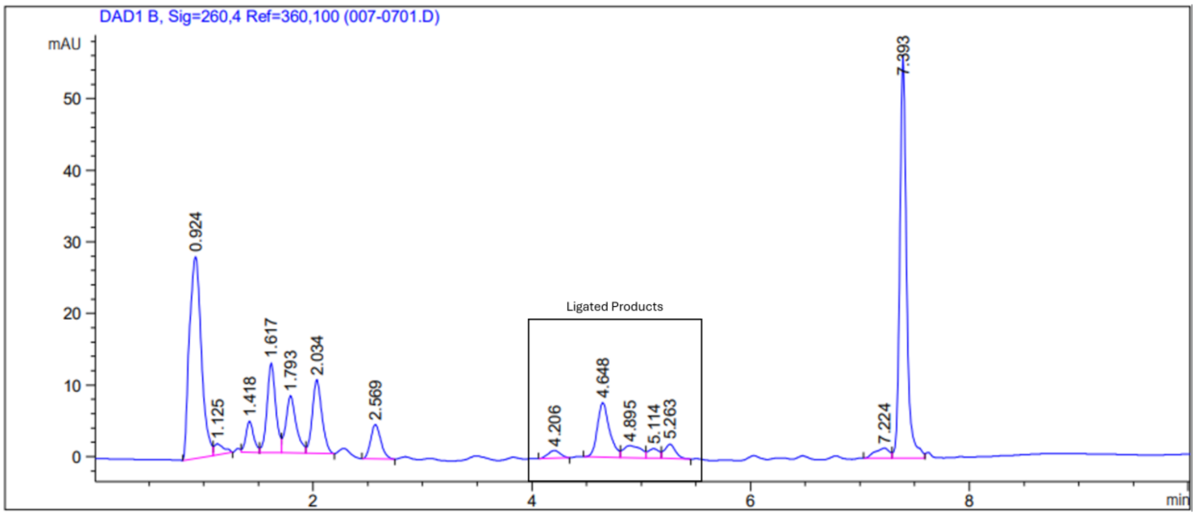


Supplementary Figure S1: HPLC Chromatograms of attempted ligations with mutated catalytic 4-base motif. A: Original E47 DNAzyme. B: 3′ T modified to an A. C: 3′ T modified to a C.

**S2A.**

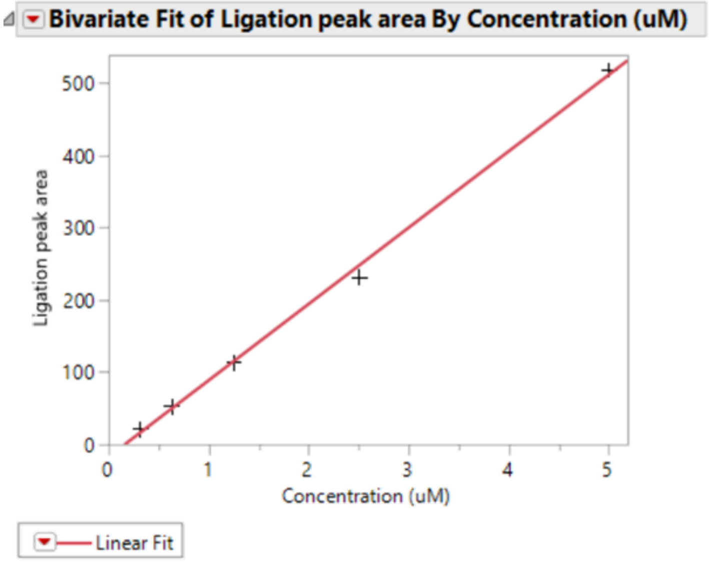


**Linear Fit**

Ligation peak area = -17.09732 + 105.75389*Concentration (uM)

**Summary of Fit**

| | |
|---|---|
| RSquare | 0.997831 |
| RSquare Adj | 0.997108 |

**S2B**

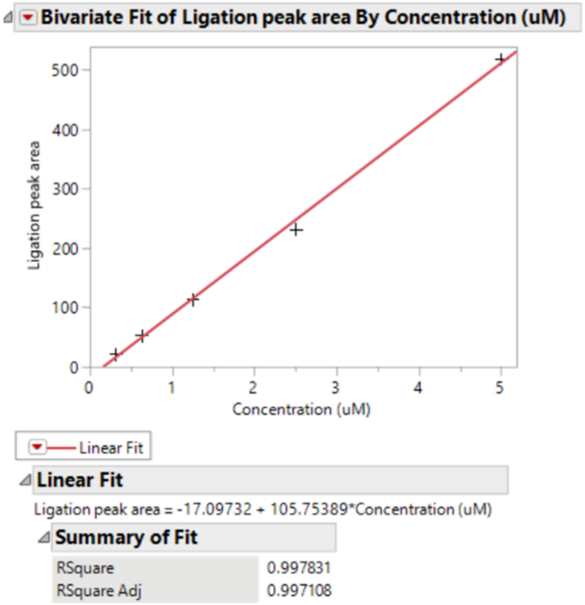


**S2C**

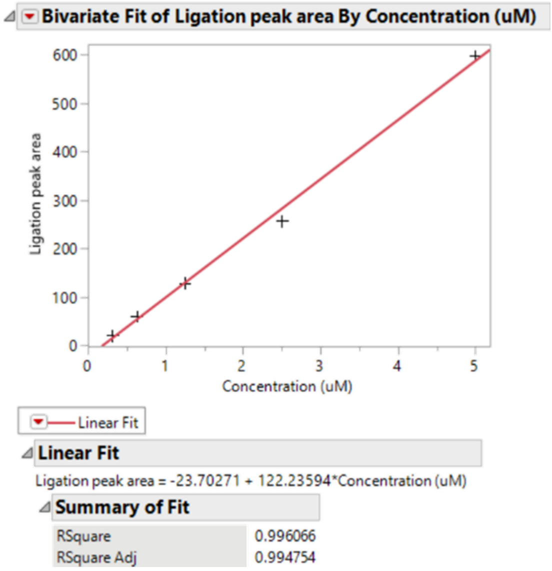


Supplementary Figure S2: Standard curves for HPLC quantification of ligation reaction yield. S2A: Standard curve for main text figure 3A. S2B: Standard curve for main text figure 3B. S2C: Standard curve for main text figure 3C.

| Sequence name | Sequence (5'-3') |
| --- | --- |
| E47 Original Catalyst | CGGATAGTGTTCTTTCGCTAGACCATGTGACGCATGGTGAGATGCTT |
| E47 original S2 | GGAACACTATCCG |
| E47 original S1 P | AAGCATCTCAAGC/3Phos/ |
| E47 Catalyst C1modG | CGGATAGTGTTCTTTGGCTAGACCATGTGACGCATGGTGAGATGCTT |
| E47 S2 C1modG | CGAACACTATCCG |
| E47 Catalyst C1modT | CGGATAGTGTTCTTTTGCTAGACCATGTGACGCATGGTGAGATGCTT |
| E47 S2 C1modT | AGAACACTATCCG |
| E47 Catalyst G2modC | CGGATAGTGTTCTTTCCCTAGACCATGTGACGCATGGTGAGATGCTT |
| E47 S1P G2modC | AAGCATCTCAAGG/3Phos/ |
| E47 Catalyst G2modA | CGGATAGTGTTCTTTCACTAGACCATGTGACGCATGGTGAGATGCTT |
| E47 S1P G2modA | AAGCATCTCAAGT/3Phos/ |
| E47 Catalyst C3modG | CGGATAGTGTTCTTTCGGTAGACCATGTGACGCATGGTGAGATGCTT |
| E47 S1P C3modG | AAGCATCTCAACC/3Phos/ |
| E47 Catalyst C3modT | CGGATAGTGTTCTTTCGTTAGACCATGTGACGCATGGTGAGATGCTT |
| E47 S1P C3modT | AAGCATCTCAAAC/3Phos/ |
| E47 Catalyst T4modA | CGGATAGTGTTCTTTCGCAAGACCATGTGACGCATGGTGAGATGCTT |
| E47 S1P T4modA | AAGCATCTCATGC/3Phos/ |
| E47 Catalyst T4modC | CGGATAGTGTTCTTTCGCCAGACCATGTGACGCATGGTGAGATGCTT |
| E47 S1P T4modC | AAGCATCTCAGGC/3Phos/ |
| Symbol A | GGTAGGAGTTCACTGAGGATAGCAGGAAGCCTAGTATCTCAAGC/3Phos/ |
| Symbol B | GGTAGGAGTTCACTGAGCAGTACATGAAGCCTAGTATCTCAAGC/3Phos/ |
| Symbol C | GGTAGGAGTTCACTGAGTAAGTCTCGAAGCCTAGTATCTCAAGC/3Phos/ |
| Primer binding site forward | GGTAGACAAGTGACCATTGACATTCTGAGTCCAGC/3Phos/ |
| Right linker 1 | GGAACACTATCTGTATGGATCTAAGAGTCTTTCAGC/3Phos/ |
| Left linker 2 | GGATACAATGATAGAGGTTGACATTCTGAGTCCAGC/3Phos/ |
| Right linker 2 | GGTTTCGTGGTAGTCATTTGACATTCTGAGTCCAGC/3Phos/ |
| Left linker 3 | GGAACACTATCTGTATGATTTATCCGTTCTGTGAGC/3Phos/ |
| Right linker 3 | GGGACAGTTCTCAATCTTTGACATTCTGAGTCCAGC/3Phos/ |
| Left linker 4 | GGAACACTATCTGTATGTGAGGATTCACGTAACAGC/3Phos/ |
| Right linker 4 | GCAGAGATTATGTGTAGTTGACATTCTGAGTCCAGC/3Phos/ |
| Left linker 5 | GGAACACTATCTGTATGTGATTGCCCTCTTCTTAGC/3Phos/ |
| Primer binding site reverse | GGAACACTATCTGTATGCCTGATAACGATCAAGTTGT |
| Left linker attachment DNAzyme | CATACAGATAGTGTTCTTTCGCTAGACCATGTGACGCATGGTGAGATACTAGGCTTC |
| Right linker attachment DNAzyme | CTCAGTGAACTCCTACTTTCGCTAGACCATGTGACGCATGGGGACTCAGAATGTCAA |
| Linker 1 to 2 DNAzyme | AGATTGAGAACTGTCCTTTCGCTAGACCATGTGACGCATGGGAAAGACTCTTAGATC |
| Linker 2 to 3 DNAzyme | CTACACATAATCTCTGTTTCGCTAGACCATGTGACGCATGGCACAGAACGGATAAAT |
| Linker 3 to 4 DNAzyme | CCTCTATCATTGTATCTTTCGCTAGACCATGTGACGCATGGGTTACGTGAATCCTCA |
| Linker 4 to 5 DNAzyme | ATGACTACCACGAAACTTTCGCTAGACCATGTGACGCATGGAAGAAGAGGGCAATCA |
| Linker 2 to 1 DNAzyme | CGATGGTCACTTGTCTAATTTCGCTAGACCATGTGACGCATGGCACAGAACGGATAAAT |
| Linker 1 to 4 DNAzyme | CCTCTATCATTGTATCTTTCGCTAGACCATGTGACGCATGGGAAAGACTCTTAGATC |
| Linker 4 to 3 DNAzyme | CTACACATAATCTCTGTTTCGCTAGACCATGTGACGCATGGAAGAAGAGGGCAATCA |
| Linker 3 to 5 DNAzyme | ATGACTACCACGAAACTTTCGCTAGACCATGTGACGCATGGGTTACGTGAATCCTCA |
| 3-piece size marker 1 | GGTAGACAAGTGACCATTGACATTCTGAGTCCAGCGGTAGGAGTT<br>CACTGAGGATAGCAGGAAGCCTAGCATCTCAAGCGGAACACTATCTGTATGGATCTA<br>AGAGTCTTTCAGC/3Phos/ |
| 3-piece size marker 2 | GCAGAGATTATGTGTAGTTGACATTCTGAGTCCAGCGGTAGGAGTT<br>CACTGAGTAAGTCTCGAAGCCTAGTATCTCAAGCGGAACACTATCTGTATGTGAGGA<br>TTCACGTAACAGC/3Phos/ |
| 5-piece size marker | GTTAGACAAGTGACCATCGTTGACATTCTGAGTCCAGCGGTAGGA<br>GTTCACTGAGGATAGCAGGAAGCCTAGCATCTCAAGCGGAACACTATCTGTATGGAT<br>CTAAGAGTCTTTCAGCGGGACAGTTCTCAATCTTTGACATTCTGAGTCCAGCGGTAGG<br>AGTTCACTGAGCAGTACATGAAGCCTAGTATCTCAAGCGGAACACTATCTGTATGATT<br>TATCCGTTCTGTGAGCGCAGAGATTATGTGTAGTTGACATTCTGAGTCCAGCGGTAG<br>GAGTTCACTGAGTAAGTCTCGAAGCCTAGTATCTCAAGCGGAACACTATCTGTATGT<br>GAGGATTCACGTAACAGCGGATACAATGATAGAGGTTGACATTCTGAGTCCAGCGG<br>TAGGAGTTCACTGAGCAGTACATGAAGCCTAGTATCTCAAGCGGAACACTATCTGTA<br>TGTGATTGCCCTCTTCTTAGCGGTTTCGTGGTAGTCATTTGACATTCTGAGTCCAGCG<br>GTAGGAGTTCACTGAGGATAGCAGGAAGCCTAGTATCTCAAGCGGAACACTATCTGT<br>ATGCCTGATAACGATCAAGTTGT |

Supplementary information Table 1: DNA sequences used

Seagate Internal
